# Can small reductions in Rubisco content improve nitrogen use in wheat without negatively impacting biomass or grain yield?

**DOI:** 10.64898/2026.02.24.702546

**Authors:** Saqer S. Alotaibi, Jack S. A. Matthews, Steven M. Driever, Caroline A. Sparks, Martin A.J. Parry, Tracy Lawson, Christine A. Raines

## Abstract

In this study, the level of Rubisco protein was reduced in wheat using RNAi, to test the hypothesis that photosynthesis, growth, and grain yield could be maintained whilst improving nitrogen use efficiency. The RNAi Rubisco wheat plants, with a Rubisco activity of less than 70% of wild type (WT) plant levels, had reduced photosynthesis, reductions in leaf and stem biomass and decreased seed yield. Interestingly, in the wheat RNAi Rubisco lines that had a small (<30%) reduction in Rubisco activity, the seed number, total seed weight and harvest index were comparable to that of WT type plants. However, no improvement in photosynthetic nitrogen use efficiency (PNUE) was evident in any of the RNAi Rubisco lines. Notably, PNUE was lower than for WT wheat plants in the RNAi lines with more than a 30% reduction in Rubisco activity. This result was unexpected and caused by an accumulation of N in both the leaves and seeds. At present we do not have an explanation for this but one possible hypothesis is that it could be due to slower growth caused by a reduction in source strength in the RNAi plants, which in turn resulted in changes to carbon and nitrogen allocation.

**Highlight:** Wheat RNAi plants with small reductions in the amount and activity of Rubisco had a similar biomass and total seed weight to that of untransformed controls but no improvement in nitrogen use efficiency was evident.

## INTRODUCTION

An improvement in crop yield is an essential requirement to meet the future demands on food and fuel of an increasing world population (FAO, 2012). However, it is unlikely that more land can be made available for the production of food, highlighting the need to improve yields from land already designated for arable use. Yield increases per unit area of land, though highly successful in the latter half of the twentieth century, have been underpinned by high inputs of nitrogen (N) fertilizers. For example, total N fertilizers used globally in agriculture increased by more than 22% from 2005 to 2016, to over 110 million tonnes (FAOSTAT, January 2020). N fertilizers are not only expensive, but are also detrimental to the environment, highlighting the importance of good nutrient management. Improving nitrogen use efficiency (NUE) in crop plants would provide both an environmental benefit and a cost reduction, noting that a 1% increase in NUE is estimated to have a potential saving of up to $1.1 billion annually in the US alone (Kant et al., 2011). However, N plays a crucial role in achieving high levels of plant productivity and in maintaining the quantity and quality of the grain. Therefore, it will be important to ensure that any strategy to alter NUE does not have negative impacts on crop yield or quality.

Nitrogen is a major component of the photosynthetic apparatus, including the enzymes in the chloroplast stroma and the light harvesting pigments in the thylakoid membranes (Bassi et al., 2018). The enzyme ribulose-1,5-bisphosphate carboxylase/oxygenase (Rubisco) catalyses the fixation of atmospheric CO_2_ in the first step in the Calvin-Benson-Bassham cycle. Rubisco has two features that result in it being a major nitrogen investment for the plant. Firstly, Rubisco is as an inefficient enzyme with a slow catalytic rate and, secondly it catalyses two competitive reactions: carboxylation and oxygenation of ribulose-1, 5 bisphosphate (RuBP) which results in the processes of photosynthesis and photorespiration respectively (Tcherkez, Farquhar et al. 2006, Andersson 2008, Hermida-Carrera, Kapralov et al. 2016). Due to these features, the Rubisco enzyme is a major limiting factor in determining the maximum rate of CO_2_ assimilation (Evans, 1989) and is needed in high amounts in the plant. As a consequence of this Rubisco is central to the nitrogen economy of a plant (Millard, 1988; Evans, 1989; Makino, 2011), accounting for approximately 50% of total leaf protein. It has also been proposed that Rubisco acts as a nitrogen store in crop plants, contributing as much as 30% of total leaf nitrogen (Masle, Hudson et al. 1993, Stitt & Schulze 1994, Evans and Poorter 2001, Carmo-Silva, Scales et al. 2015; Hermeda-Carrera et al. 2016,).

Data obtained from plants grown in elevated CO_2_ conditions, together with modelling studies, provide evidence that under some environmental conditions, there is an over-investment in Rubisco protein (Ainsworth & Long, 2005; Stitt & Schulze, 1994). This was supported by analysis of transgenic plants with reduced levels of Rubisco, that suggested there was up to a five-fold over-investment in Rubisco protein in plants grown at lower light levels (Hudson et al., 1992; Stitt & Schulze, 1994). The implications of this is that in the UK, where crops such as wheat can frequently be exposed to moderate light levels (750 μmol m^-2^ s^-1^), the amount of Rubisco protein may not be limiting for photosynthetic carbon fixation (Stitt & Quick, 1991). This raises the hypothesis that there is an opportunity to reduce total plant N requirement by reducing Rubisco protein levels, which could help increase productivity in poor soils without needing large amounts of N fertiliser (An et al., 2018). While this strategy would be advantageous in current temperate climate conditions, it also has the potential to be of particular benefit for crops growing under future elevated atmospheric CO_2_ concentrations, where carboxylation would be increased relative to oxygenation, reducing further the amount of Rubisco needed under current CO_2_ conditions.

Wheat is an important crop worldwide and it has been proposed that there is a potential over-investment of leaf N in Rubisco in wheat. Therefore a small reduction in the amount of Rubisco enzyme per unit leaf area could represent an opportunity to reduce crop N requirements and improve NUE, whilst sustaining wheat yield and reduced application of N fertilizer inputs (Mitchell, Grant et al. 2000). The Rubisco holoenzyme is composed of eight large subunits (LSu) and eight small subunits (SSu). The LSu is encoded by a single chloroplast gene, *rbcL* but the SSu is encoded in the nucleus by a family of *rbcS* genes. The levels of the holoenzyme are known to be controlled by the availability of the SSu and hence it is possible to change the total amount of the Rubisco enzyme by altering the level of the SSu. To explore the impact of reduced levels of Rubisco protein on photosynthesis, growth and grain yield in wheat plants, grown in glasshouse conditions, we have used an interference RNA (RNAi) construct targeted to the *rbcS* gene family to produce transgenic wheat plants with a range of reduced levels of the Rubisco protein and Rubisco activity.

## MATERIALS AND METHODS

### Construction of RNAi vector and production of transgenic plants

A fragment of DNA (345 bp), encoding the most highly conserved region of the wheat SSu protein coding sequence, was used to construct the *RbcS* RNAi cassette. This fragment of DNA was cloned in both the sense and antisense orientation at the 5’ and 3’ ends, as described in Travella et al, (2006), with the addition of EcoRV and XmaI restriction sites at the outermost borders. The DNA was synthesised (Invitrogen, Inchinnan, Scotland, UK) and cloned into the plasmid vector at the DraIII sites to generate a level 0 plasmid. The functional RNAi construct (Fig. 1a) (WRubiscoRNAi) was produced by digesting the level 0 plasmid with EcoRV and XmaI restriction enzymes, and cloning the resulting fragment into the corresponding sites of an expression vector (pRRes14.101; Rothamsted Research, West Common, Harpenden, UK) which placed the RNAi cassette under the control of a maize ubiquitin promoter containing an intron. This construct also carried a cassette containing the *bar* gene under the control of the maize ubiquitin promoter and intron. The *bar gene* encodes resistance to glufosinate ammonium-containing herbicides e.g. Basta, which allows selection of transformed plants in tissue culture using phosphinothricin (the active ingredient of glufosinate ammonium herbicides). The recombinant *RbcS* RNAi plasmid was introduced into wheat embryos (cv. Cadenza) by particle bombardment, as described by Sparks and Jones (2009).

**Figure 1.**
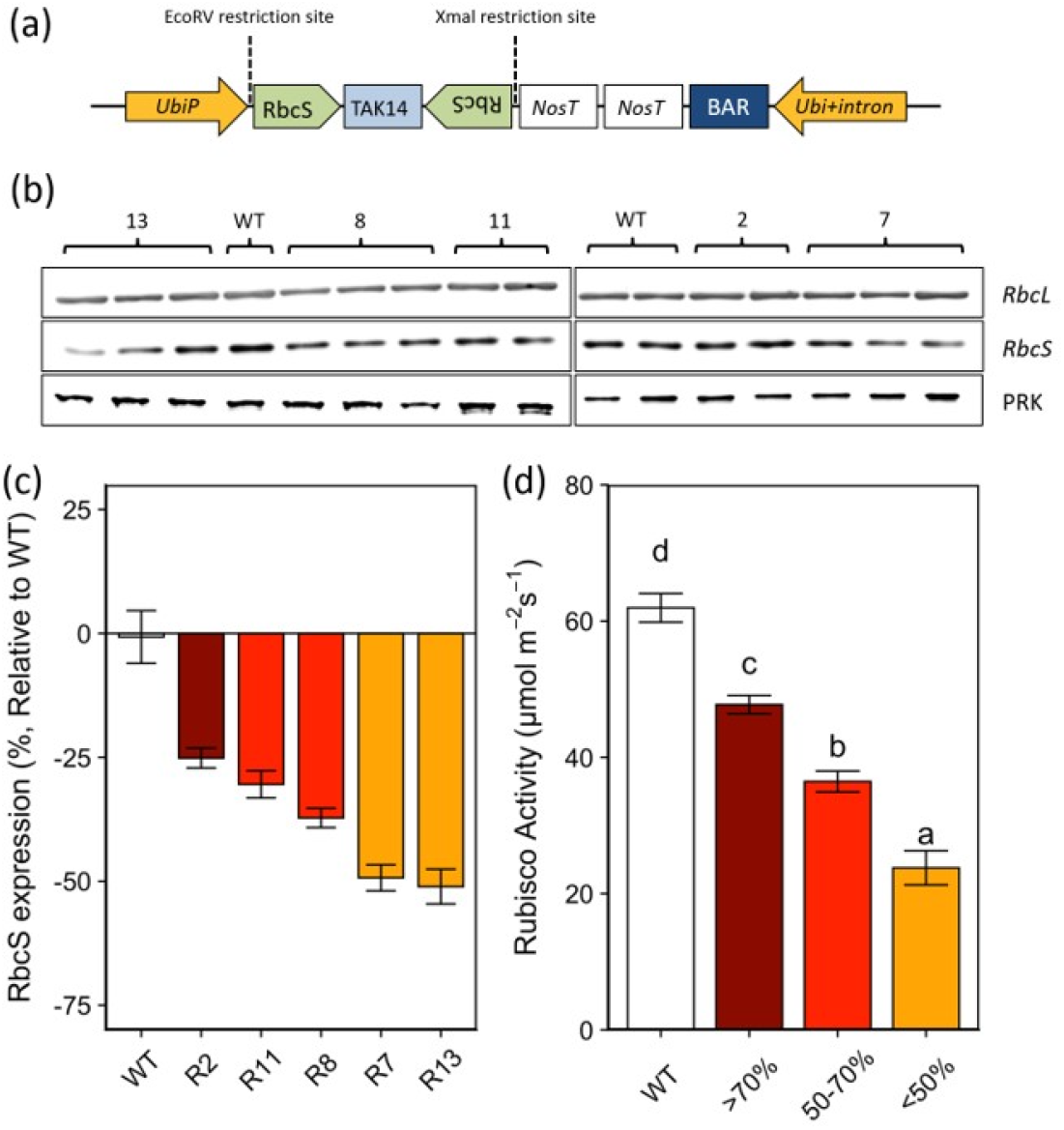
Production and analysis of transgenic wheat with reduced levels of Rubisco enzyme. (a) Schematic of the *rbcS* RNAi construct used for transformation of wheat. *UbiP*, maize ubiquitin-1 promoter; Ubi+ intron, maize ubiquitin-1 promoter plus intron; TAK14 (b) Immunoblot analysis of leaf protein extracts from transgenic T_2_ wheat lines using Rubisco holoenzyme (Foyer *et al*., 1993), and PRK (phosphoribulokinase)antibodies ((Klein and Salvucci 1995). (c) *rbcS* gene expression, determined using qRT-PCR in transgenic T_2_ wheat lines. (d) Plants are grouped based on Rubisco activity > 70% (R2), between 50-70% (R11 & R8) and < 50% (R7 & R13) of WT Rubsico activity. Measurements were carried out on fully developed flag leaves (Zadoks 4.1-4.5). Error bars represent mean +/-SE, n = 4-6. Asterisks indicate significant difference relative to WT plants (p<0.05). The means for the groups of plants were compared using ANOVA with a Tukey post-hoc test.

From two separate bombardment events, 13 independent T_0_ *rbcS* RNAi plants were identified by PCR and quantitative reverse transcriptase PCR (qPCR), and allowed to self-fertilize. The resulting T_1_ plants were screened using qPCR and western blot analysis, and five lines were selected and allowed to self fertilise, producing T_2_ plants that were used in this study.

### Growth conditions

Transgenic wheat plants were produced as described above and the T2 generation seed germinated and grown in Levington F2+S compost (Fisons, Ipswich, UK) in a climate controlled growth chamber for three weeks (22°C, 60% relative humidity, 16 h photoperiod). Selected seedlings were then transferred into 4 L pots and grown in a semi-controlled environment glasshouse (25-32°C day/18°C night) with a 16 h photoperiod provided by supplementing natural irradiance with sodium lamps to a minimum light level of 350 μmol m^-2^ s^-1^ photosynthetically active radiation (PAR). All plants were well watered and regularly moved to minimize the effect of spatial and temporal variation in growth conditions.

### DNA extraction from wheat leaves

DNA was extracted from one-week old wheat leaves as follows: tissues were placed into 2 ml micro-tubes, frozen on dry ice and freeze-dried overnight. Freeze-dried tissue was ground according to the manufacturers instructions using a Qiagen grinder (Retsch mill, tissueLyser (type MM300), Qiagen, Manchester, UK,) and 600 μl of extraction buffer (0.1 M Tris-HCl, 0.05 M EDTA, 1.25% SDS, pre-heated to 65°C) added and incubated at 65°C for 30 min. Samples were cooled down to 4°C for 15 min before adding 300 μl of 6 M ammonium acetate, left to stand for 15 min before centrifugation at 13000 xg for 15 min at 4°C to collect the precipitated proteins and plant tissues. The supernatant (600 μl) was recovered in a tube containing 360 μl of isopropanol and the DNA pelleted by centrifugation at 13000 xg for 15 min at 4°C. The DNA pellets were washed using 70% ethanol (v/v) and centrifuged at 13000 xg for 15 min at 4°C, removing the supernatant and allowing to air dry. Pellets were re-suspended in 50 μl of purified water, and stored at -20°C for future analysis.

### Amplification of specific gene fragment for screening

To confirm the presence of the introduced DNA in the transgenic wheat plants, PCR was used. The reactions were prepared as follows: 1 μL of forward primer (10 pmol), 1 μL of 10 pmol reverse primer (Table 1), 2 μL of 10 x buffer (containing 0.1 M Tris-HCl; pH 8.3, 0.1 mg ml^-1^ gelatine, 500 mM KCl, and 25 mM MgCl_2_), 0.2 μL of Dream *Taq* polymerase enzyme and 0.2 μL of dNTPs were mixed with 1.5 μL of extracted DNA sample and 14.1 μL sterile, double purified water. The PCR specifications were as follows: 3 min at 98°C, followed by 35 cycles of 30 s each at 98°C, 30 s at 60-68°C (based on primers Tm for each gene), 1 min at 72°C, and finally, 10 min at 72°C and stored at 4°C.

**Table 1:**
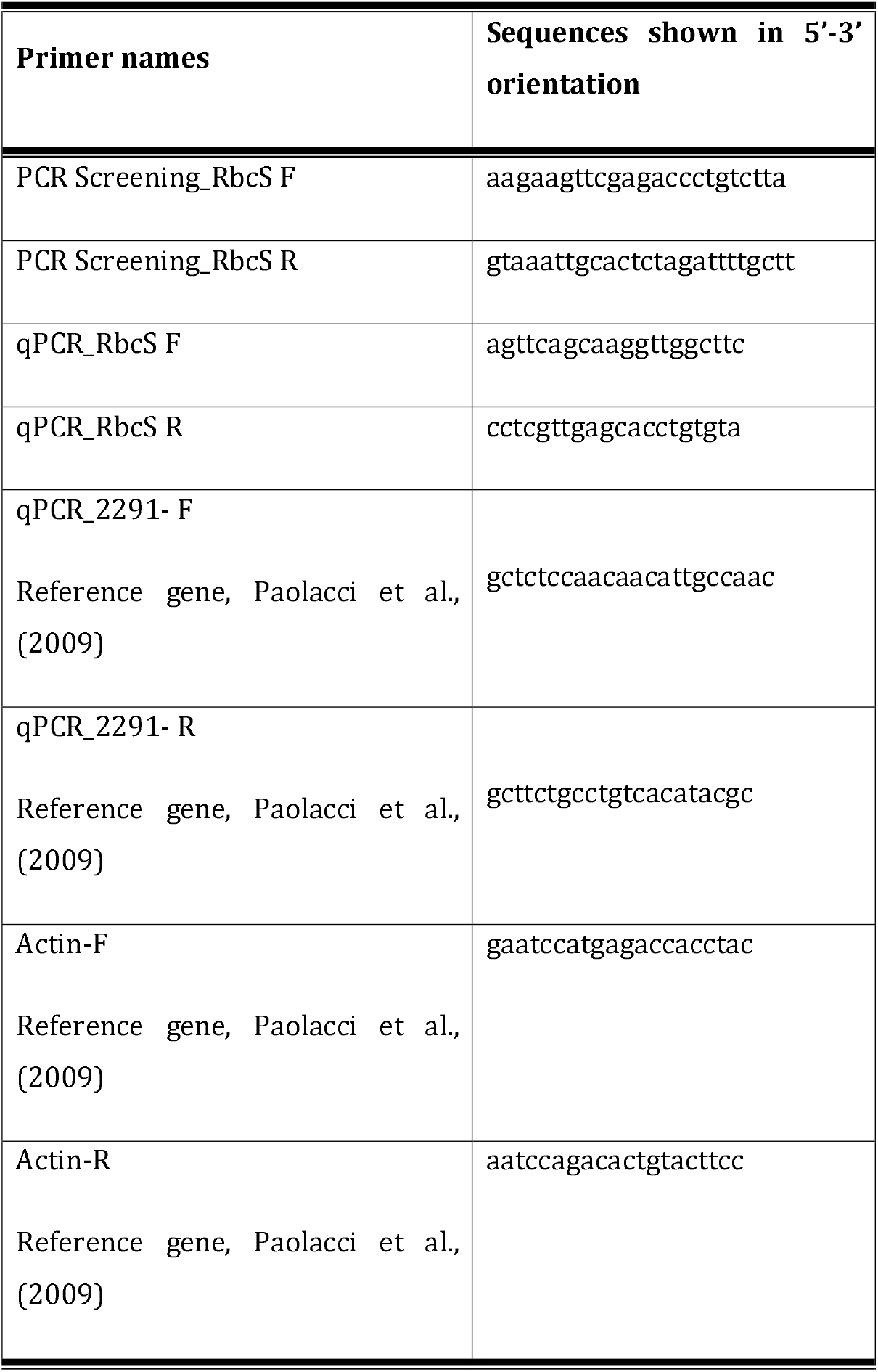
Primers were designed to identify the introduced Rubisco sequence and used to screen for transgenic Rubsico RNAi lines using qPCR.

### RNA extraction from wheat leaves and cDNA synthesis

Total RNA was isolated from frozen leaf material (*ca* 0.1 g fresh weight; FW) ground in liquid nitrogen using Tri-reagent (Sigma, Welwyn Garden City, UK), modified from Hilario and Mackay (Driever, Simkin et al, 2017) with the following additional steps: 3 M sodium acetate (30 µl) and 750 µl of ice-cold absolute ethanol were added and centrifuged at 13 000 xg at 4°C for 15 min and the sample was then left on dry ice for 20 min. The RNA pellets were washed with 75% ethanol and centrifuged for 5 min. Finally, the pellets were allowed to air-dry for 10 min before re-suspension in 50 µl of purified water and stored at −80°C. The extracted RNA was quantified using a NanoDrop spectrophotometer (Nanodrop Products, Thermofisher, Inchinnan, UK). cDNA was synthesized from mRNA using SuperScript first strand cDNA synthesis (Invitrogen, Thermofisher, Inchinnan, UK) as specified by the manufacturer. The cDNA was stored at -20°C prior to qPCR analysis.

### qPCR analysis to determine the abundance of the introduced rbcS gene

Synthesized cDNA was amplified using qPCR to determine if the transformed genes were expressed and to provide information on the abundance of the transcripts. A mix containing 7.5 µl of Sybre No-ROX mix (Bioline Reagents Ltd, London, UK) plus 10 pmol of forward primer and reverse primer was (1 μL of each) (Table 1) was added to the master mix (9 μL) then in triplicate into 96 well plates including cDNAs (6 μL) and mixed together by pulse centrifugation. qPCR cycling conditions were as follows: 95°C for 3 min, (95°C for 10 s, 62°C for 30 s) (45 cycles). qPCR reactions were performed using SensiFast SYBR No-ROX mix (Bioline Reagents Ltd, London, UK) as specified by the manufacturer.

### Protein extraction and quantification

Leaf samples (*ca* 0.2 g FW) were ground in liquid nitrogen, protein extracted and quantified essentially as described by (Harrison, Willingham et al. 1998). Equal total protein (10 µg) was loaded onto 12% (w/v) Tris-SDS-PAGE gel (Bio-rad, Watford, UK), separated and transferred onto a cellulose nitrate membrane (0.45 µm pore, Millipore Immobilon-P Transfer Membrane)(GE Healthcare Life Science, Munich, Germany). Proteins were detected using antibodies raised against the wheat Rubisco holoenzyme Rubisco (Foyer *et al*., 1993), phosphoribulokinase (PRK; Klein and Salvucci, 1995), and detected using horseradish peroxidase conjugated to the secondary antibody and Pierce ECL chemiluminescence detection reagent (Thermo Scientific, Rockford, IL, US) and visualized using a FUSION FX chemiluminescence detection system (PEQLAB Ltd, Southampton, UK).

### Total Rubisco activity

The activity of Rubisco was measured as previously described by Pengelly et al., (2010), with some modifications. Total protein was extracted by grinding 3 cm^2^ of frozen wheat leaf tissue in liquid N_2_ and 800 µl of protein extraction buffer (50 mM HEPES (pH 8.2), 5 mM MgCl_2_, 1 mM EDTA, 10% Glycerol, 0.1% Triton X-100, 2 mM Benzamidine, 2 mM Aminocaproic acid, 10 mM DTT and 0.5 mM PMSF and 10 µl of protease inhibitor cocktail (Sigma) add and mixed. Samples were transferred to tubes and spun at 13000 xg for 2 min at 4°C and 200 µl of the supernatant was transferred into fresh tubes and used directly for assays. For the Rubisco activity assay, 2.5 µl of leaf extract was combined with 242.5 µl of assay buffer (50 mM EPPS-NaOH pH=8, 10 mM MgCl_2_, 0.5 mM EDTA, 1 mM ATP, 5 mM phosphocreatine, 20 mM NaHCO_3_, 0.2 mM NADH, 50 U ml^-1^ creatine phosphokinase, 0.2 mg carbonic anhydrase, 50 U ml^-1^ 3 phosphoglycerate kinase, 40 U ml^-1^ glyceraldehyde 3-phosphate dehydrogenase, 113 U m^-1^ triose phosphate isomerase, 39 U ml^-1^ glycerol 3-phosphate dehydrogenase) followed by reaction initiation by addition of 5 µl of 21.9 mM of RuBP substrate. The activity of Rubisco enzyme was calculated from the reduction of NADH absorbance at 340 nm using a spectrophotometric micro-plate reader (SpectroStar Omega, BMC Labtech, Aylesbury, UK).

### Measurement of photosynthetic efficiency using *A/C*_*i*_ (net photosynthetic rate/intercellular CO_2_ concentration) response curves

Gas exchange was measured using a Li-Cor 6400XT portable gas exchange system, with an integral blue-red LED light source (Li-Cor, Lincoln, Nebraska, US). For all gas exchange measurements, a constant flow rate was set at 300 µmol s^-1^, with cuvette conditions maintained at an ambient CO_2_ concentration of 400 µmol mol^-1^ and a leaf temperature of 25°C with a vapour pressure deficit of 0.9 kPA. All measurements were performed on a fully expanded flag leaf, Zadoks stage 4.1-4.5. The response of net CO_2_ assimilation rate (*A*) to intercellular CO_2_ concentration (*C*_*i*_) was measured at a saturating light intensity of 2000 μmol m^-2^ s^-1^. Leaves were initially stabilized at ambient CO_2_ concentration of 400 µmol mol^-1^ and a measurement taken, before the CO_2_ concentration was decreased to 300, 200, 100 and 50 µmol mol^-1^, following which readings were taken at the initial value of 400, and at 650, 900, 1200, 1500 µmol mol^-1^. The maximum velocity of Rubisco for carboxylation (*Vc*_*max*_), the maximum rate of electron transport demand for RuBP regeneration (*J*_max_), were derived by curve fitting as described by Sharkey et al. (2007) using the Rubisco kinetic constants for wheat (Carmo-Silva et al., 2010). Operational assimilation rate under ambient conditions (*A*400) and maximal carboxylation rate (*A*_*max*_) were determined from assimilation values recorded at 400 and 1500 μmol mol^−1^ CO_2_ concentration, respectively.

### Analysis of growth, biomass, and yield

Stages of wheat growth and development were measured using the Zadoks scale (Zadoks et al., 1974), with leaves and tillers counted every week to record changes in growth and development. Biomass data was collected when all plants reached physiological maturity (Zadoks growth stage 9.1-9.2). Leaves, stems, ears and seeds were harvested, counted, and dried at 70°C until a constant weight was obtained, then final dry weights of each were determined. Ears were threshed and the number of seeds counted and seed weight determined. The fraction of total N in leaves was analysed using an element analyser based on the micro-Dumas combustion method.

### Statistical Analyses

Statistical Analysis was conducted using R software (www.r-project.org; version 3.6.3). Where more than one group existed, one-way ANOVA was used to test for single factor differences. When significant differences were observed, Tukey post-hoc comparisons of group. Means were used to compare the different treatments.

## RESULTS

### Production and identification of *rbcS* RNAi wheat plants

An *rbcS* RNAi construct, containing 345 bp of DNA from the protein coding region of the Ssu sequence, was designed in both the antisense and sense orientation at the 5’ and 3’ ends respectively, and separated by an intron derived from the wheat TAK14 gene as described in Travella et al., (2006) with the addition of EcoRV and XmaI restriction sites (Fig. 1a). This construct was introduced into wheat (cadenza) using particle bombardment and a number of primary transformants (T0) were produced, from which 13 independent transformants carrying our gene of interest were identified by PCR analysis. These plants were allowed to self-fertilize and the resulting T_1_ seed collected. The resulting T_1_ plants were screened for expression of the *rbcS* transcript and western blotting used to determine changes in the levels of the Rubisco protein (data not shown). Based on this analysis five T_1_ lines (R2, 11, 8, 7 & 13) were taken through to the T_2_ generation. The T_2_ plants were grown under semi-controlled conditions in a glasshouse. Western blot analysis of total protein extracts from the *rbcs* RNAi plants showed that the levels of the Rubisco SSu were reduced in all of selected lines and there was no decrease in the levels of the Calvin cycle enzyme PRKase (PRK) or Rubisco large subunit (RbcL) (Fig. 1b). Quantification of *rbcS* cDNA from the five independent lines and WT plants showed reduced expression levels in all RNAi lines compared to WT plants (Fig. 1c), with reductions in expression levels ranging from 25-52%. Total extractable Rubisco enzyme activity revealed transgenic lines with a range of Rubisco activities that were grouped into plants with > 70% (R2), between 50-70% (R11 & R8) and < 50% (R7 & R13) of WT Rubsico activity (Fig. 1d).

### Effect of reduced Rubisco activity on photosynthesis and photosynthetic capacity

To investigate the effect of the decreased level of Rubisco activity on photosynthesis, net CO_2_ assimilation rates (*A*) were measured as a function of intercellular [CO_2_] (*A*/*C*_*i*_) in the flag leaf of WT and RNAi Rubisco lines, and data grouped according to the amount of Rubisco activity in the plant (Fig. 2). In the flag leaf the impact of reduction in Rubisco activity was similar in all of the RNAi lines with reductions in the light saturated reate of photosynthesis observed in all of the plants. The light saturated and CO_2_ saturated rate of A (A1500 was significantly lower in all lines with <70% Rubisco activity. The maximum rate of carboxylation by Rubisco (*V*_cmax_) determined from the *A*/*C*_i_ curves, although lower in all lines, was not signficnatly different. Whilst the maximum electron transport demand for RuBP regeneration (*J*_max_) was signifcnalty lower in Rubisco RNAi plants, with <70% Rubisco (Table 2).

**Table 2.** Rubisco^Act^, *A*_400_, *A*_1500_, *V*_*cmax*_ and *J*_*max*_ values calculated from A/Ci curves. The maximum rate of Rubisco enzyme for carboxylation (*V*_cmax_), the maximum rate of electron transport require for RuBP regeneration (*J*_max_), the ambient CO_2_ rate of photosynthesis (A_sat_) and the CO_2_-saturated rate of photosynthesis (Amax) were solved by curve fitting as described by equations described in Sharkey et al., (2007).

| | Rubisco<sup>Act</sup> | $A_{400}$ | $A_{1500}$ | $V_{cmax}$ | $J_{max}$ |
| --- | --- | --- | --- | --- | --- |
| WT | 63.2±2.43<sup>a</sup> | 29.9±0.58<sup>a</sup> | 49.9±1.24<sup>a</sup> | 145.8±9.25<sup>a</sup> | 213.0±8.82<sup>a</sup> |
| >70% | 44.3±2.00<sup>b</sup> | 25.6±0.95<sup>b</sup> | 38.5±2.18<sup>a</sup> | 142.8±5.33<sup>a</sup> | 206.4±13.6<sup>a</sup> |
| 50-70% | 37.7±8.52<sup>c</sup> | 23.5±1.38<sup>b</sup> | 37.0±0.92<sup>b</sup> | 123.4±3.77<sup>a</sup> | 200.4±4.18<sup>b</sup> |
| <50% | 30.2±1.95<sup>d</sup> | 21.7±2.18<sup>b</sup> | 36.4±0.92<sup>b</sup> | 121.8±12.29<sup>a</sup> | 196.4±6.69<sup>b</sup> |

**Figure 2.**
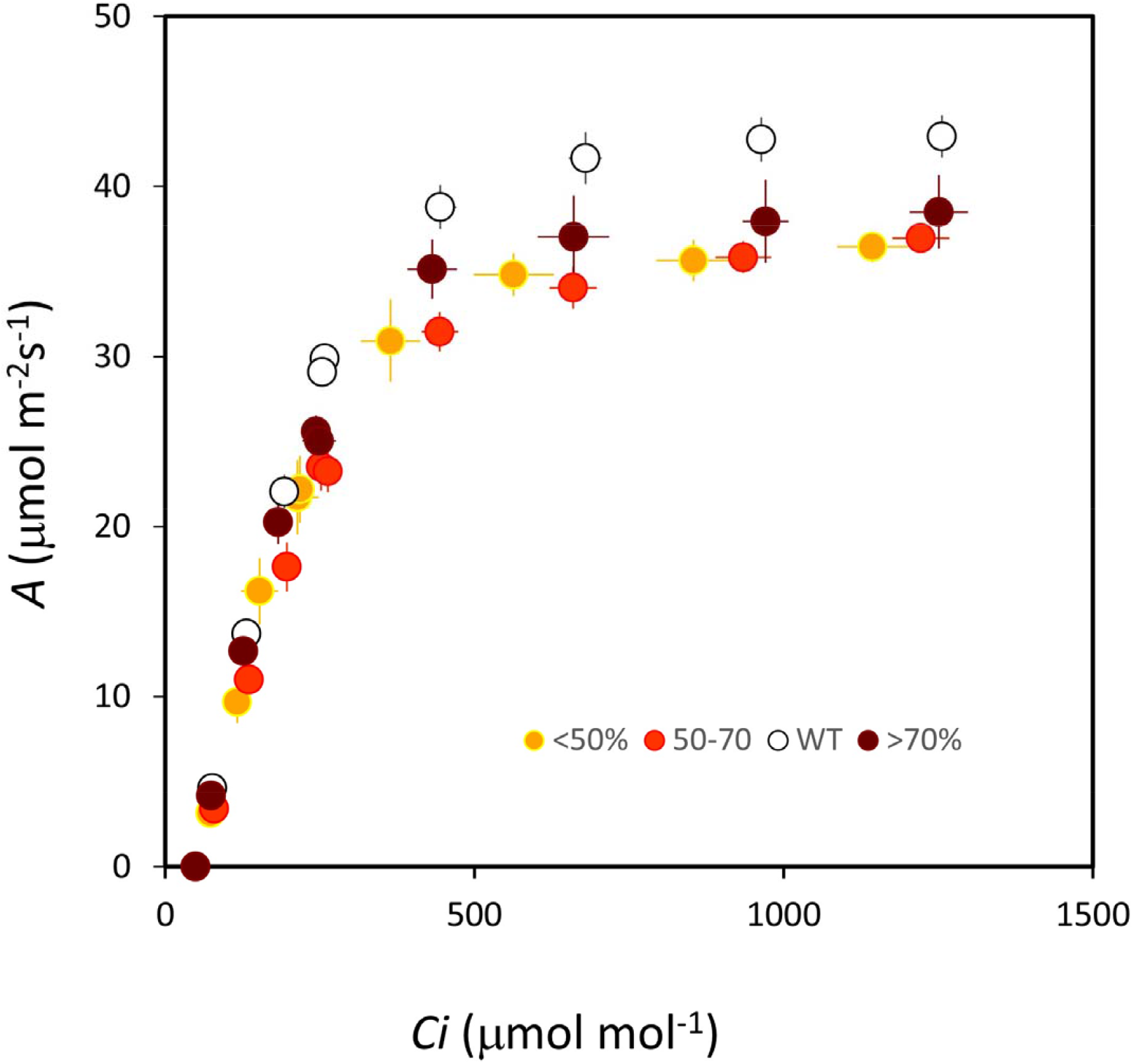
Effect of reductions in Rubisco activity on whole leaf photosynthesis. CO_2_ assimilation as a function of of intercellular CO_2_ concentration (*C*_*i*_). Plants are grouped based on Rubisco activity and measurements were carried out on fully developed flag leaves (Zadoks 4.1-4.5). Error bars represent mean +/-SE, n = 4-6.

### Growth and biomass of Rubisco RNAi plants

To determine the impact of reduced Rubisco activity on plant growth and development the RNAi Rubisco and WT plants were grown in a semi-controlled environment glasshouse and Zadoks stage of development tracked weekly. In the RNAi plants with the smallest reductions in Rubisco (>70% WT) growth was similar to that of WT plants (Fig. 3a). In contrast the RNAi Rubisco lines with the greatest reduction in Rubisco activity (<50% WT) showed a slower growth. As a result the RNAi lines R7, R8 and R13 were shorter (Fig. 3b) and produced less biomass than WT plants at Zadoks stage 7.0. This reduction in the rate of growth continued until Zadoks stage 9.1-9.2, shown in (Fig. 3b), when the plants were harvested.

**Figure 3.**
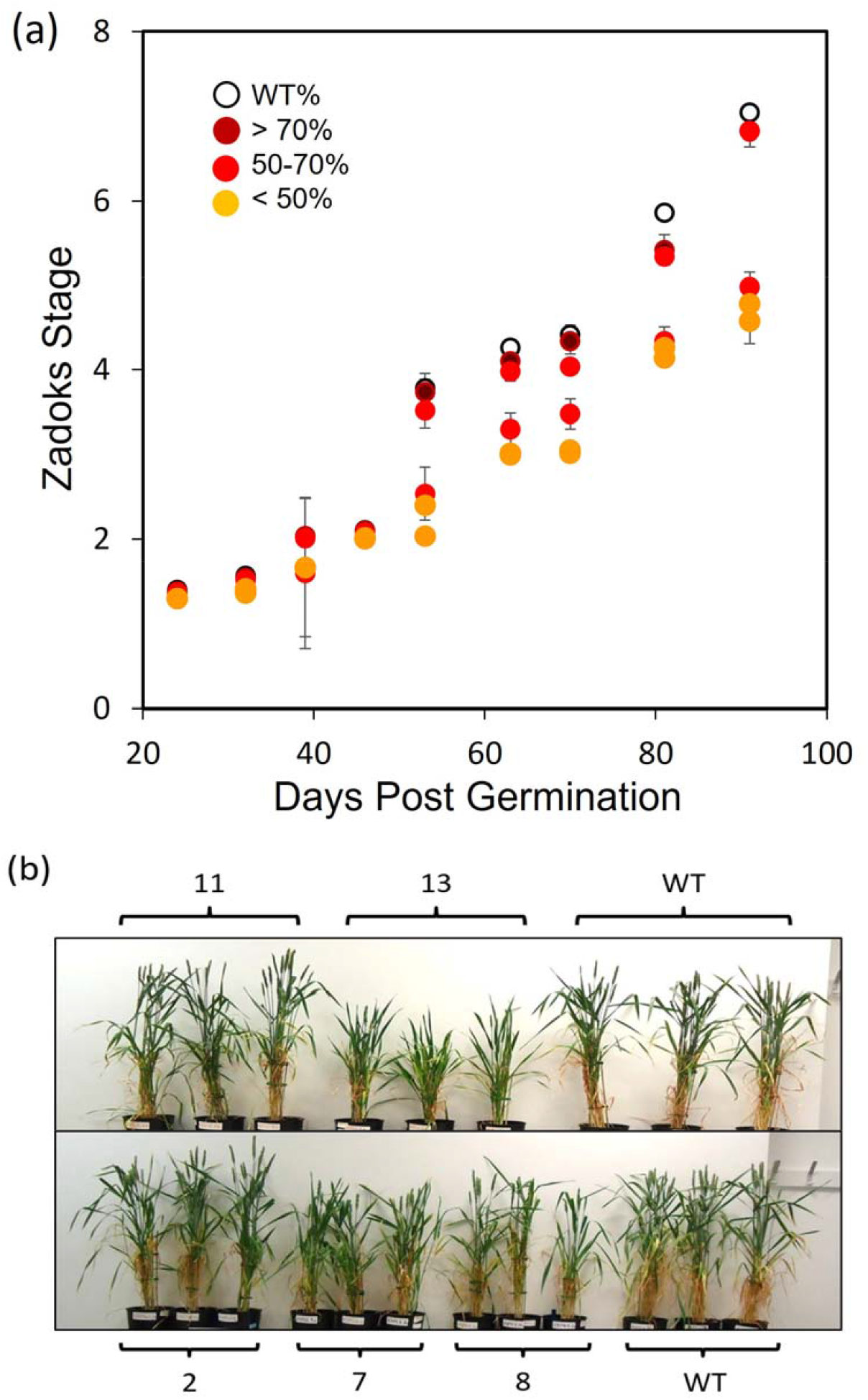
Developmental and growth characteristics of the SSu RNAi wheat plants. (a) Time course of Zadok stage, measured weekly until Zadoks 7.0. (b) Representative phenotypes of wheat transgenic T_2_ lines with different Rubisco activities relative to WT plants at Zadoks 9.1-9.2. Plants are grouped based on Rubisco activity > 70% (R2), between 50-70% (R11 & R8) and < 50% (R7 & R13) of WT Rubsico activity.Error bars represent mean +/-SE. n = 4-6.

Rubisco activity was closely correlated with overall plant growth, decreases in plant stem, leaf and overall biomass observed with decreasing activity (Fig. 4 a-c). The Rubisco RNAi plants with the largest reductions in Rubisco activity produced the lowest amount of total vegetative biomass compared to the WT plants (Fig. 4a). This was due to a reduction in both the leaf and stem biomass (Fig. 4b and 4c). The plants with the smallest reductions in Rubisco activity had a similar stem and leaf biomass to that of the WT plants. No strong correlation in the leaf/stem ratio was observed in response to a decrease in Rubisco activity (Fig. 4d).

**Figure 4.**
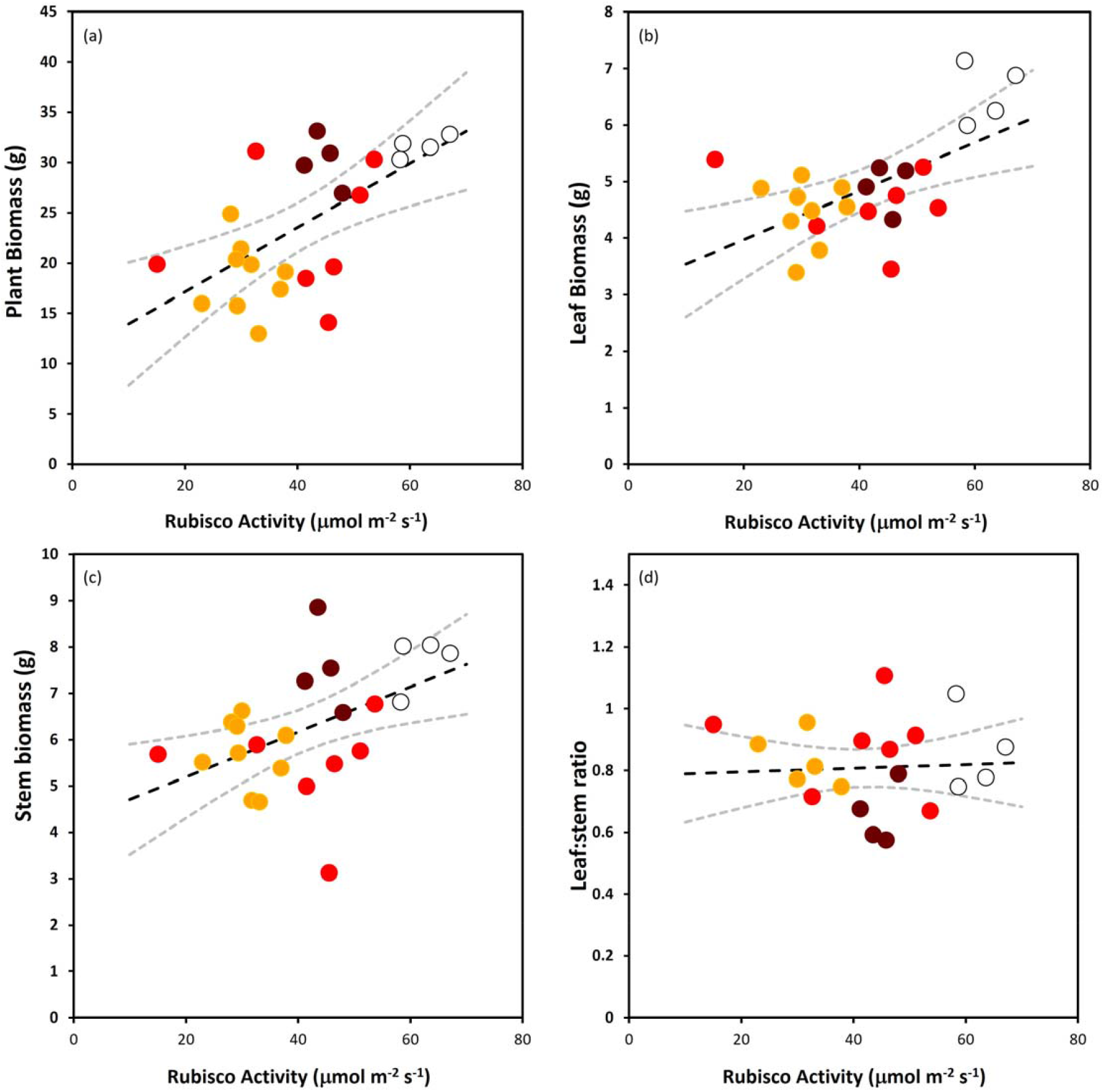
Vegetative biomass of WT and SSu RNAi plants correlated to Rubisco activity. (a) Total above ground plant biomass; (b) Total leaf biomass; (c) Stem biomass; (d) Leaf/stem ratio. Plants were harvested at full maturation Zadoks 9.1-9.2. Each symbol represents a single plant. The regression line with a 95% confidence interval (grey dashed lines) was generated in Excel using regression and calculating confidence limits

Significant changes in the biomass and numbers of reproductive parts of the RNAi plants were recorded. The number of ears per plant decreased even in plants with small reductions in Rubisco activity and, in the lines with the greatest reductions, this was found to be close to half of that in WT plants (Fig. 5a). Seed number per ear was also lower in the RNAi Rubisco plants and, in those plants with less than 70% WT Rubisco activity seed weight was below 50% of WT plants (Fig. 5b). Unexpectedly, weight per seed was increased significantly in all of the RNAi Rubisco plants (Fig. 5c). In RNAi Rubisco lines with less than 50% of WT activity a substantial reduction in the total number of seed per plant was observed (Fig. 5d), driven by a combination of a decrease in ears per plant (Fig. 5a) and seeds per ear (Fig. 5b). This resulted in a significant reduction in total seed weight in those lines with less than 70% Rubisco relative to WT plants (Fig. 5e). In contrast plants with Rubisco activity greater than 70% of WT levels, had a similar total seed weight to that of the WT plants. In these plants the small decrease in total seed number was compensated for by the increase in individual seed weight (Fig. 5c). The harvest index (the ratio of seed yield to above ground vegetative biomass) was reduced in RNAi Rubisco plants but a statistically significant impact was seen in only those plants with less than 50% Rubisco activity compared to WT (Fig. 5f).

**Figure 5.**
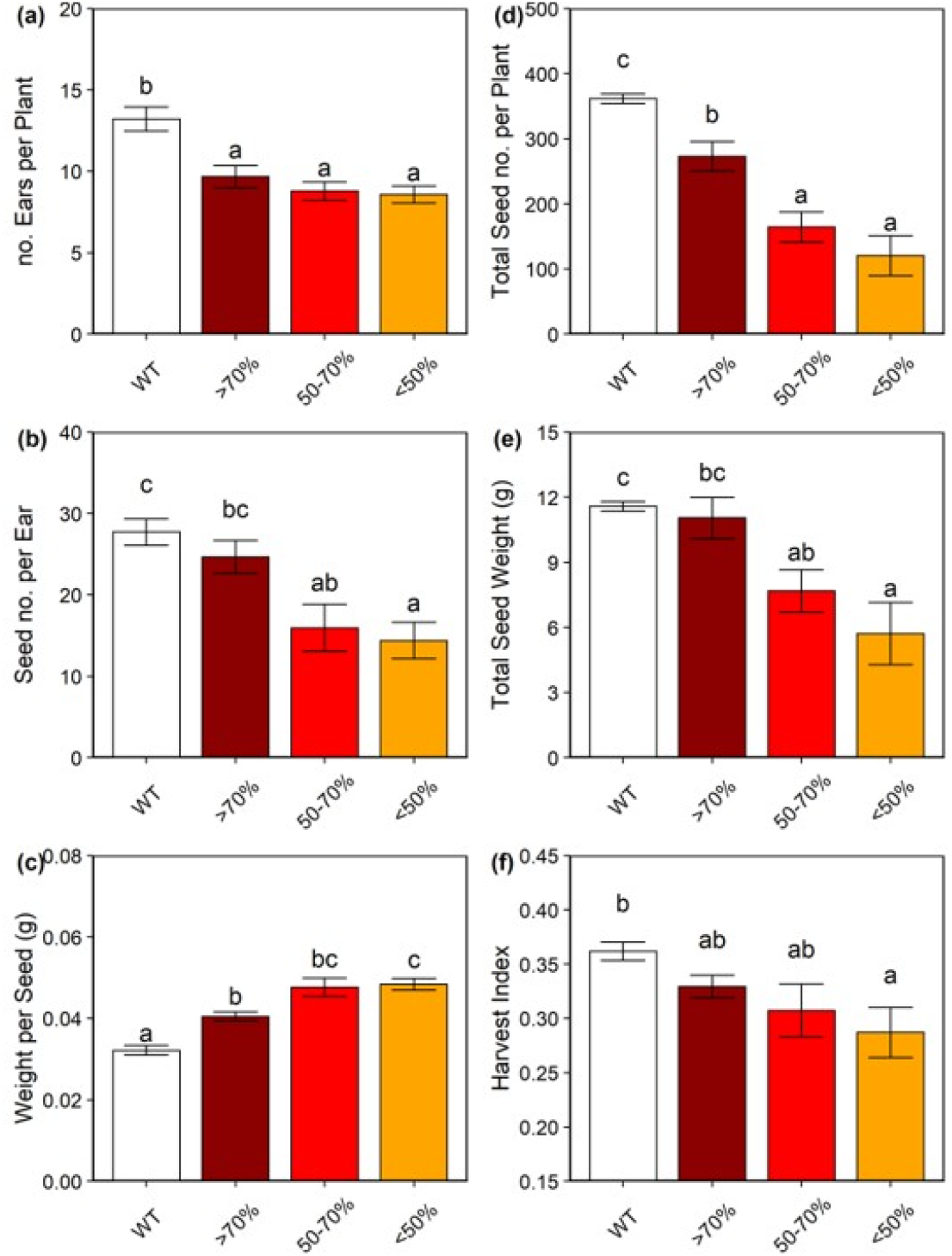
Seed production and weight of wheat SSu RNAi plants harvested at Zadoks 9.1-9.2. (a) Number of ears per plant; (b) Seed number per ear; (c) Weight per seed; (d) Total seed number per plant; (e) Total seed weight; (f) Harvest index. Plants are grouped based on Rubisco activity and were harvested at full maturation Zadoks 9.1-9.2. Error bars represent mean +/-SE. n = 4-6. Asterisks indicate significant difference to WT (p<0.05).

### Effect of reduced Rubisco activity on nitrogen content and harvestable yield

To determine the levels of nitrogen in the leaves and seeds of the Rubisco RNAi and WT plants, plants were harvested and analysed when they reached Zadoks stage 9.1-9.2. It was found that plants with the lowest level of Rubisco activity (those below 50% of WT activity) had significantly higher levels of leaf and seed nitrogen compared to the WT plants (Fig. 6a and b), with a ca. 100% increase in leaf nitrogen content and a ca. 30% increase in seed nitrogen. In contrast, RNAi Rubisco plants, with activity above 70% of WT activity, exhibited nitrogen contents comparable to WT plants. As a result of the large increases in nitrogen content, PNUE determined as the ratio of CO_2_ assimilation to leaf organic nitrogen content, was significantly reduced in the RNAi Rubisco plants with the lowest activity of Rubisco at both ambient (PNUE_sat_, Fig. 6c) and saturating CO_2_ (PNUE_max_, Fig. 6d). The RNAi plants with Rubisco activities above 70% of WT, had no significant difference in values of PNUE_sat_ and plants with > 70% activity had PNUE_max_ values similar to WT.

**Figure 6.**
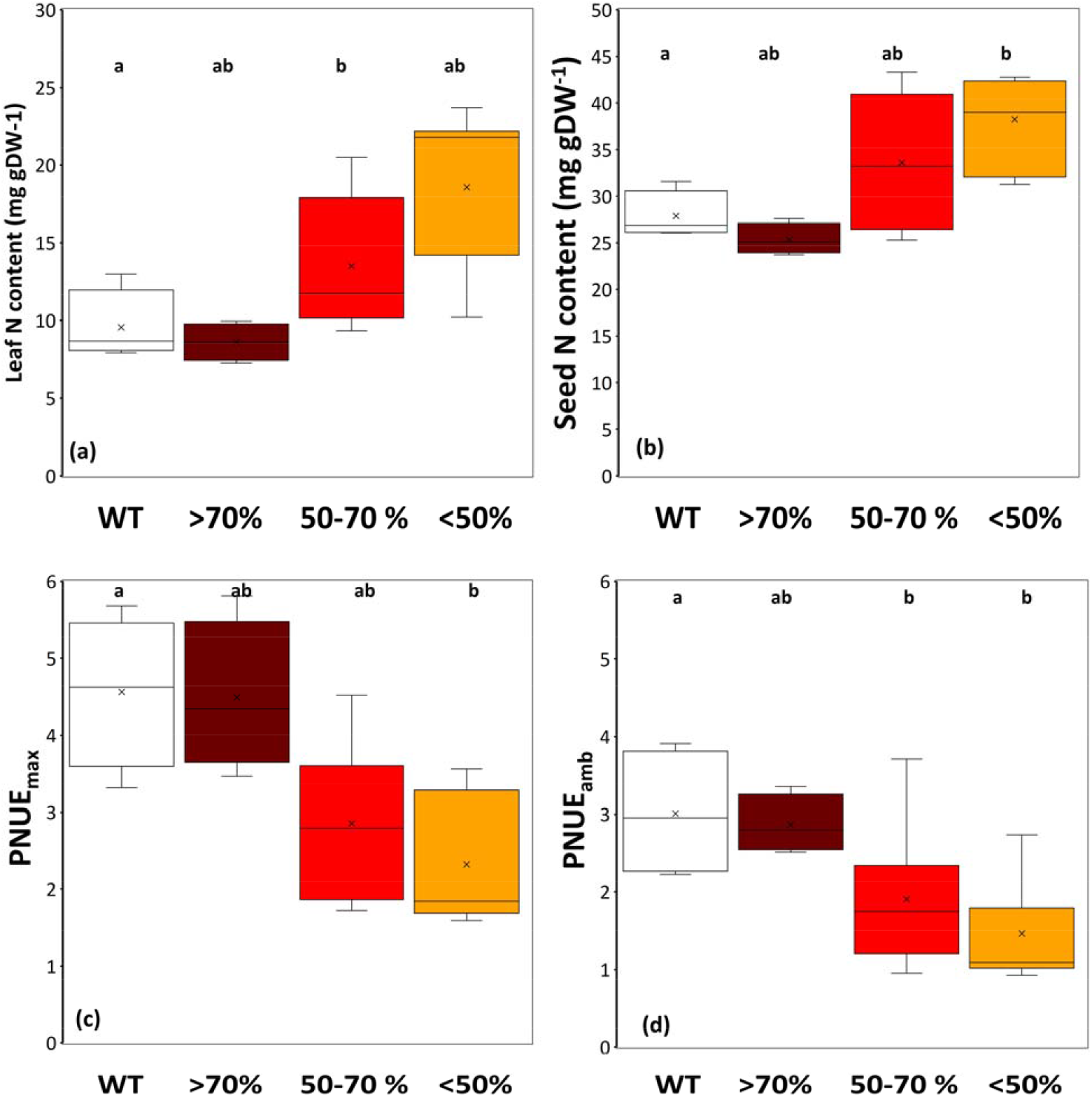
Nitrogen content and photosynthetic nitrogen use efficiency of WT and SSu RNAi plants. (a) Leaf nitrogen content; (b) Seed nitrogen content; (gDW is mg per g dry weight) Plants are grouped based on Rubisco activity and were harvested at full maturation Zadoks 9.1-9.2. (c-d) Photosynthetic nitrogen use efficiency at ambient (c, 400ppm) and high (d, 1500ppm) CO_2_ concentrations calculated from A/Ci response curves. Box plots represent middle 50% of the data bound by the 25 and 75^th^ percentile, and bars the highest and lowest values. The X in the box represent the mean and the solid line the median.Letters indicate significant difference between groups at p<0.05.

The relationship between Rubisco activity, nitrogen content, seed weight and harvest index is shown in Figure 7. The N content of both the leaves and seeds increased with decreasing Rubisco activity and plants with the lowest Rubico activity accumulated more N in both the leaves and the seed than WT plants (Fig. 7a and 7b). In contrast, total seed weight per plant was decreased with decreasing Rubisco activity (Fig. 7c). Harvest index also displayed a positive relationship with Rubisco activity, with the lowest HI evident in plants with the lowest Rubisco activity (Fig. 7d).

**Figure 7.**
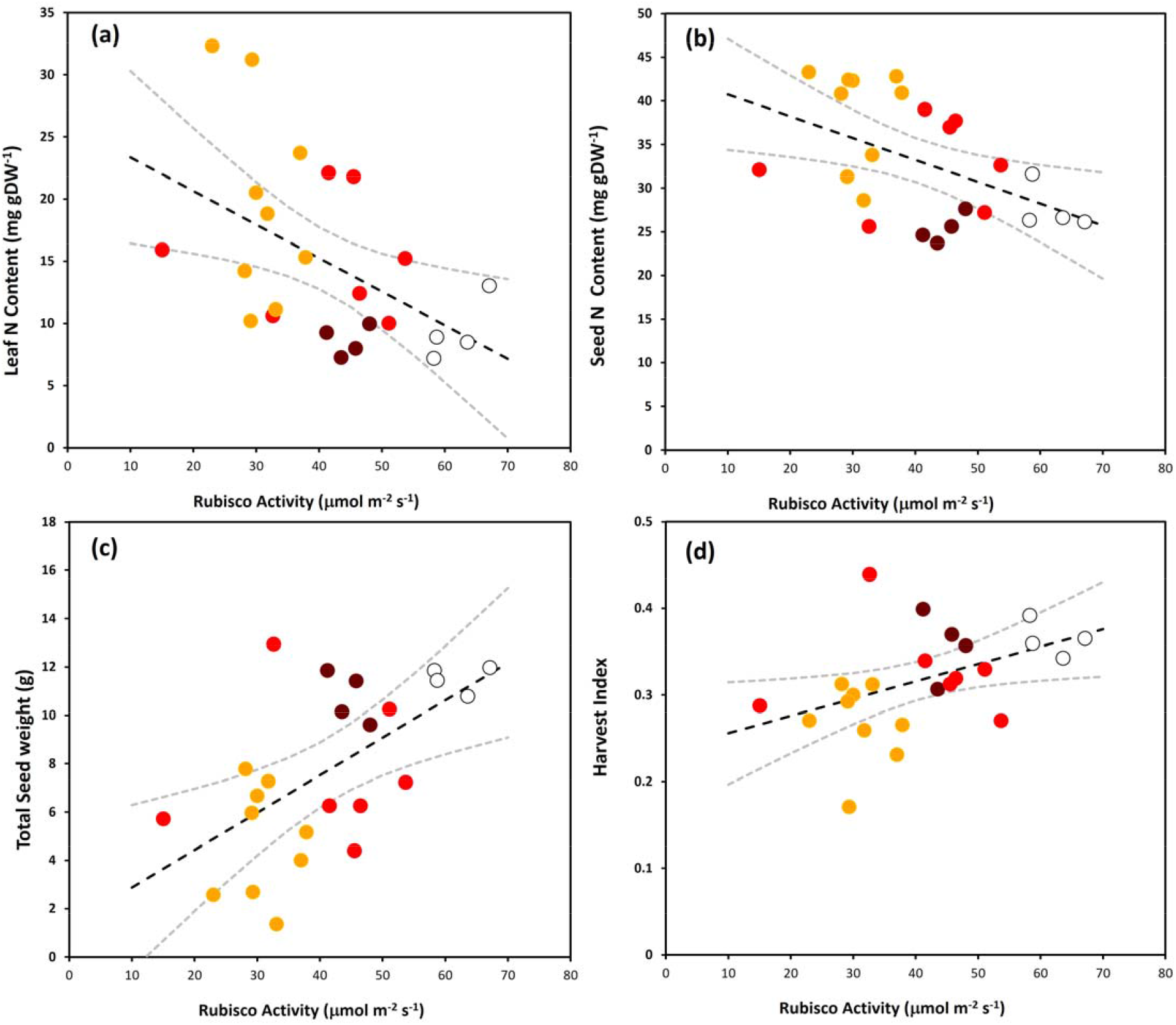
Nitrogen content, seed weight and harvest index in WT and SSu RNAi plants correlated to Rubisco activity. (a) Leaf nitrogen content; (b) Seed nitrogen content; (c) Total seed weight; (d) Harvest index. Symbols indicate individual plants; WT (black symbols)). Plants were harvested at Zadoks stages 9.1-9.2. The regression line with a 95% confidence interval (grey dahsed lines) was generated in Excel using regression and calculating confidence limits

## DISCUSSION

In this study, the level of Rubisco protein and activity in wheat was reduced using a SSu RNAi construct, to test the hypothesis that photosynthesis, growth, and grain yield could be maintained whilst improving nitrogen use efficiency. In agreement with previous studies, RNAi Rubisco plants with reductions in Rubisco content had a reduced photosynthetic performance, growth and biomass. Interestingly, the wheat RNAi Rubisco plants that had Rubisco activity of more than 70% maintained seed number, total seed weight and harvest index similar to that of wild-type plants as well as PNUE_amb_ and PNUE_max._ Plants with <70% Rubisco activity had reduced PNUE_amb_ and PNUE_max_ levels compared with WT. These results were unexpected and at present we do not have a definitive explanation for the observed decrease in PNUE, which was caused by an increase in N content. However, our results would suggest that it is likely to be due to a reduction in source strength, resulting in changes to carbon and nitrogen allocation and reductions in growth.

In all of the RNAi wheat plants decreases in the photosynthetic rate were observed in the flag leaf, when measured at either elevated or ambient CO_2_ concentrations (as shown in the *A/C*_*i*_ curves). A reduction in the maximum rate of electron transport to sustain RuBP regeneration (*J*_*max*_) in the RNAi plants was observed, together with reduction in the maximum rate of carboxylation (*V*_*cmax*_). Considering there was no significant difference observed in the photosynthetic rates between the wheat Rubsico RNAi plants and the WT plants, the reason for the differential reduction in biomass was not immediately evident. One explanation could be that the effect of the RNAi construct was stronger during earlier stages of development. Support for this explanation can be seen in the time course of development (Fig. 3a) which shows that effects on growth in the plants with the lowest levels of Rubisco protein were evident at ∼ 40-50 days post germination and that these plants never attained the size or biomass of WT plants. A similar effect was observed in rice antisense Rubisco plants, where it was proposed that a stronger expression of the introduced construct early in development, led to slower growth of these plants resulting in a negative impact on the final biomass (Kanno et al, 2017).

Differential changes were evident in growth parameters, seed production and biomass between plants Rubisco activity of more than 70% and those with less than 70%, relative to WT plants. The differences between these two groups of Rubisco RNAi plants was highly significant in terms of time to reach a specific Zadok stage in development, total biomass, total seed weight and seed number per plant. This agrees with previous studies in both tobacco and rice that have shown that reductions in Rubisco protein of greater than 40% relative to WT resulted in slower growth, fewer leaves, stems and ears, as well as delayed development and flowering time (Jiang & Rodermel, 1995, Tsai et al., 1995). In some cases these changes have been linked to delays in the early stages of development (Masle et al., 1993; Makino, 2003) but in other studies growth was reduced significantly without any change in the time to reach a given developmental stage (Makino et al., 2000; Bassi et al., 2018). Our work taken together with that of others suggests that there is a threshold of Rubisco content and activity of ca. 70% that of wild-type plants, below which plants exhibit more marked decreases in photosynthetic rate, growth, biomass production and seed yield. Interestingly, in both the rice and tobacco Rubisco antisense studies it was shown that the negative impact on growth was almost entirely cancelled out when the antisense plants were grown under elevated CO_2_ conditions (Masle et al., 1993; Makino et al., 2000). A recent study showed that when rice plants, with small reductions in Rubisco protein (10-20%) were grown in elevated CO_2_ conditions, higher rates of photosynthesis were observed and an increase in both biomass and grain was observed in comparison to WT plants (Makino 2021). This is an interesting result providing support for down regulation of Rubisco in crops to benefit growth in high CO_2_ environments, predicted to occur in the near future.

One aspect of the work reported here was to explore the possibility of reducing Rubisco protein whilst maintaining photosynthesis and PNUE. Counter intuitively, in the wheat RNAi plants, with Rubisco activity less than 70% of WT, higher levels of N accumulated, resulting in a reduction PNUE_amb_ but only in plants with less than Rubisco activity > 50% was PNUE_max_ reduced. Perhaps this result is not surprising given the slow growth phenotype and the N accumulation in this group of RNAi plants. A similar increase in N was also observed in field grown tobacco *rbcS* RNAi lines, with below 50% of WT levels (Meecham-Henshold et al., 2019). These results are in contrast to some reports that suggest a decrease in Rubisco content should lead to reductions in total nitrogen concentration and photosynthesis, as reported in rice (Makino et al, 2000; Makino, 2003; Suzuki et al., 2009, 2010; Sudo et al., 2014; Kanno et al., 2017; Suganami et al., 2018) and cotton (Guo et al., 2016). These contrasting results may be explained by species differences and or environmental conditions under which the plants were grown as the impact of Rubisco levels on photosynthesis and growth has been shown to be dependent on growth conditions (Stitt and Schulze, 1994).

The reason for the accumulation of N could be due to internal changes in sink/source balance. Evidence to support this hypothesis is the reduction in seed production observed in the wheat Rubisco RNAi plants, which was not only due to the reduction in the number of ears produced per plant, but also to a reduced number of seeds produced per ear. It has been reported previously that reductions in photosynthetic carbon assimilation and changes in N status, brought about by reductions in Rubisco protein, can play a role in altering the early stages of development (Masle et al., 1993; Fichtener et al., 1993). It has also been proposed that the changes in growth of *rbcS* RNAi rice plants may be caused by internal signals associated with the suppression of photosynthesis and/or the products related to it (Makino et al., 2000; Suzuki et al., 2009). Although it has been known for some time that changes in leaf carbon metabolism can alter development, the mechanisms underlying these responses remain to be resolved and, exploitation of transgenic plants with altered photosynthetic metabolism may provide novel insight into the regulatory processes involved (Raines and Paul, 2005; Lawson, Bryant et al, 2006).

## CONCLUSION

We have shown that plants with small decreases in Rubisco content via the RNAi suppression of the SSu, have small reduction in photosynthesis but can maintain growth, and grain yield similar to that of WT plants. However in these plants no increases in nitrogen use efficiency were observed, with levels similar to WT plants. This finding does not support the hypothesis that *rbcS* plants with small reductions in Rubisco content of less than ca. 20-30% have the potential for improvement in nitrogen use efficiency under current CO_2_ or nitrogen replete conditions. To test further the potential of reductions in Rubisco to benefit wheat production in future climatic conditions the effects of growing the wheat Rubisco RNAi plants in elevated CO_2_ with N replete and N limiting conditions should be explored. This is of relevance due to the current high demands for nitrogen fertilization in modern agriculture and the cost and environmental impacts that this imparts. Furthermore, there is a need to engineer new varieties that are able to sustain yields in low nutrient soils, and improve nitrogen use in low to moderate yielding fields. The results here also raise the possibility that combining reduction in Rubisco holoenzyme with other enzymes in the Calvin-Benson-Bassham cycle (e.g. SBPase overexpression), increases in biomass and yield could be achieved (Driever et al., 2017, Raines, 2023).

## Author contributions

SSA and SD designed the construct and the experiments. SSA selected the RNAi lines used in the analysis, acquired the data and undertook the first round of analysis and interpretation. JASM re-analysed the data, prepared the figures for the paper and drafted the manuscript. C.A.S. performed the wheat transformation and initial line selection (T_0_) and revised the manuscript. CAR and M.A.J.P. conceived the study. CAR, JASM & TL drafted the first versions of the paper and TL and CAR redrafted and finalised the manuscript.

## Data availability statement

All primary data will be available on publication and materials published in this paper are available on request to the corresponding authors at

